# VAPPER: High-throughput Variant Antigen Profiling in African trypanosomes

**DOI:** 10.1101/492074

**Authors:** Sara Silva Pereira, John Heap, Andrew R. Jones, Andrew P. Jackson

## Abstract

**Background:** Analysing variant antigen gene families on a population scale is a difficult challenge for conventional methods of read mapping and variant calling due to the great variability in sequence, copy number and genomic loci. In African trypanosomes, hemoparasites of humans and animals, this is complicated by variant antigen repertoires containing hundreds of genes subject to various degrees of sequence recombination.

**Findings:** We introduce Variant Antigen Profiler (VAPPER), a tool that allows automated analysis of variant antigen repertoires of African trypanosomes. VAPPER produces variant antigen profiles for any isolate of the veterinary pathogens *Trypanosoma congolense* and *Trypanosoma vivax* from genomic and transcriptomic sequencing data and delivers publication-ready figures that show how the queried isolate compares with a database of existing strains. VAPPER is implemented in Python. It can be installed to a local Galaxy instance from the ToolShed (https://toolshed.g2.bx.psu.edu/) or locally on a Linux platform via the command line (https://github.com/PGB-LIV/VAPPER). The documentation, requirements, examples, and test data are provided in the Github repository.

**Conclusion:** Our approach is the first to allow large-scale analysis of trypanosome variant antigens and establishes two different methodologies that may be applicable to other multi-copy gene families that are otherwise refractory to high-throughput analysis.

## Background

Advances in next-generation sequencing have enabled researchers to produce high-throughput genomic data for diverse pathogens. However, analysing multi-copy, contingency gene families remains challenging due to their abundance, high mutation and recombination rates, and unstable gene loci [1]. Yet, these gene families are often involved in many processes of pathogenesis, including antigenic variation, virulence, host use, and immune modulation in a multitude of pathogens [2–4]. A prime example of a crucial gene family lacking the necessary analytic tools for high-throughput analysis is the Variant Surface Glycoprotein (*VSG*) superfamily in African trypanosomes [5].

African trypanosomes are extracellular hemoparasites that cause human sleeping sicknessand animal African trypanosomiasis (AAT). Their genomes contain up to 2500 *VSG* genes [6] dispersed through specialized, hemizygous chromosomal regions called subtelomeres, smaller chromosomes, and less frequently in the core of megabase-sized diploid chromosomes. The *VSG* genes encode variant surface glycoproteins, GPI-anchored proteins that coat the entire surface of the parasite in the bloodstream of the mammal host, which function mostly in antigenic variation and immune-modulation [7]. Sporadically, specific *VSG* genes have been shown to evolve other functions, not related to antigenic variation, such as conferring human infectivity to *T. brucei gambiense* (*TgsGP* gene) [8, 9] and *T. brucei rhodesiense* (*SRA* gene) [10, 11], resistance to the drug suramin (*VSG^sur^* gene) [12], and mediating the transport of transferrin (*TfR* genes) [13, 14].

As they are key players in host-trypanosome interaction, understanding *VSG* diversity and its impact in pathology, disease phenotype and virulence is of foremost importance in trypanosome research [4]. However, the *VSG* repertoire cannot be accurately analysed using conventional approaches of read mapping and variant calling. Attempts to bypass this challenge have resulted in alternative approaches using manually-curated *VSG* gene databases for specific *T. brucei* strains [6, 15–17], but to the best of our knowledge there is no automated tool for the systematic analysis of *VSG* from any trypanosome genome. Thus, we have developed <u>V</u>ariant <u>A</u>ntigen <u>P</u>rofil<u>er</u> (VAPPER), a tool that examines *VSG* repertoires in DNA/RNA sequence data of the main livestock trypanosomes, *Trypanosoma congolense* and *T. vivax*, and quantifies antigenic diversity. This results in a variant antigen profile (VAP) that can be compared between isolates, locations, and experimental conditions [18]. In this paper we briefly present how VAPPER can be used to further our knowledge of antigenic diversity and variation.

## Findings

### The service

VAPPER is primarily intended for producing and comparing VAPs of livestock trypanosomes, without the need for complex bioinformatic processes. It is available online through the Galaxy ToolShed [19] for a local Galaxy server [20], and as a Linux package for local installation. The program has three pipelines, specific for each organism (*T. congolense* or *T. vivax*) and input data type (genome or transcriptome). VAPPER requires quality-filtered, trimmed, paired sequencing reads in FASTQ format [21] or assembled contigs in FASTA format [22]. Results are presented in tables of frequencies, heatmaps, and Principal Component Analysis (PCA) plots, visualized as HTML files or exported to PDF or PNG format. A typical workflow is shown in Fig. 1.

**Figure 1.**
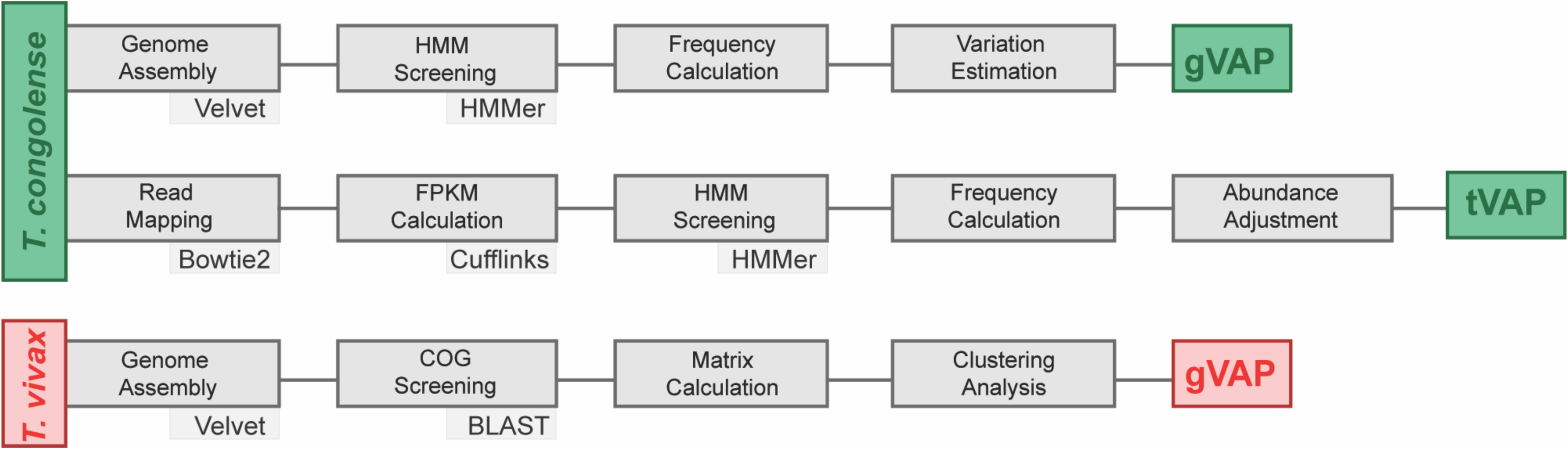
Methodological workflow according to species (*T. congolense* or *T. vivax*) and input data [genomic (gVAP) or transcriptomic (tVAP)].

For *T. congolense* genomic VAPs (gVAP), VAPPER starts with genome assembly of raw, short reads using Velvet 1.2.10 [23]. Assembled contigs are screened for pre-defined protein motifs described by a hidden Markov model using HMMER 3.1b2 [24] after 6-frame translation. A detailed description of the universal protein motifs and their biological significance in a recent manuscript [18], but, in summary, each protein motif or motif combination is diagnostic of a specific phylotype [18]; therefore, phylotype frequency can be calculated from the HMMER output. The proportions of each phylotype represent the gVAP and are recorded in a table of frequencies. The gVAP produced is also placed in the context of a *T. congolense* genome database supplied with VAPPER (N=97, [18, 25]), which is regularly updated. This is achieved through a Euclidean distance-based clustering analysis. Results are presented as two heatmaps with corresponding dendrograms, one showing phylotype frequency, and the other showing frequency deviation from the population mean. They are also shown as a PCA plot and a table of frequencies.

For *T. congolense* transcriptomic analyses (tVAP), VAPPER performs read mapping using Bowtie 2 2.2.6 [26], reference-based transcript assembly and abundance calculation using Cufflinks 2.2.1 [27], and *VSG* transcript screening and phylotype assigning as described for gVAP. The proportions of each phylotype are then adjusted for transcript abundance based on the Cufflinks output (Fig. 1). The tVAP is presented as a weighted bar chart and compared to the gVAP of the reference (Fig. 2c). Ideally, the user would provide their own reference genome for the mapping step. As that is not always possible, especially for field isolate analysis, we provide two reference genomes, the IL3000 Kenyan isolate [14, 28], and the Tc1/148 Nigerian isolate [29, 30]. Choosing the most adequate reference for the sample being analysed may potentially improve the VAPPER results by increasing mapping sensitivity. However, we have previously shown that closely related *T. congolense* strains (i.e. with short genetic distances) do not always have equally related *VSG* repertoires [18].

**Figure 2.**
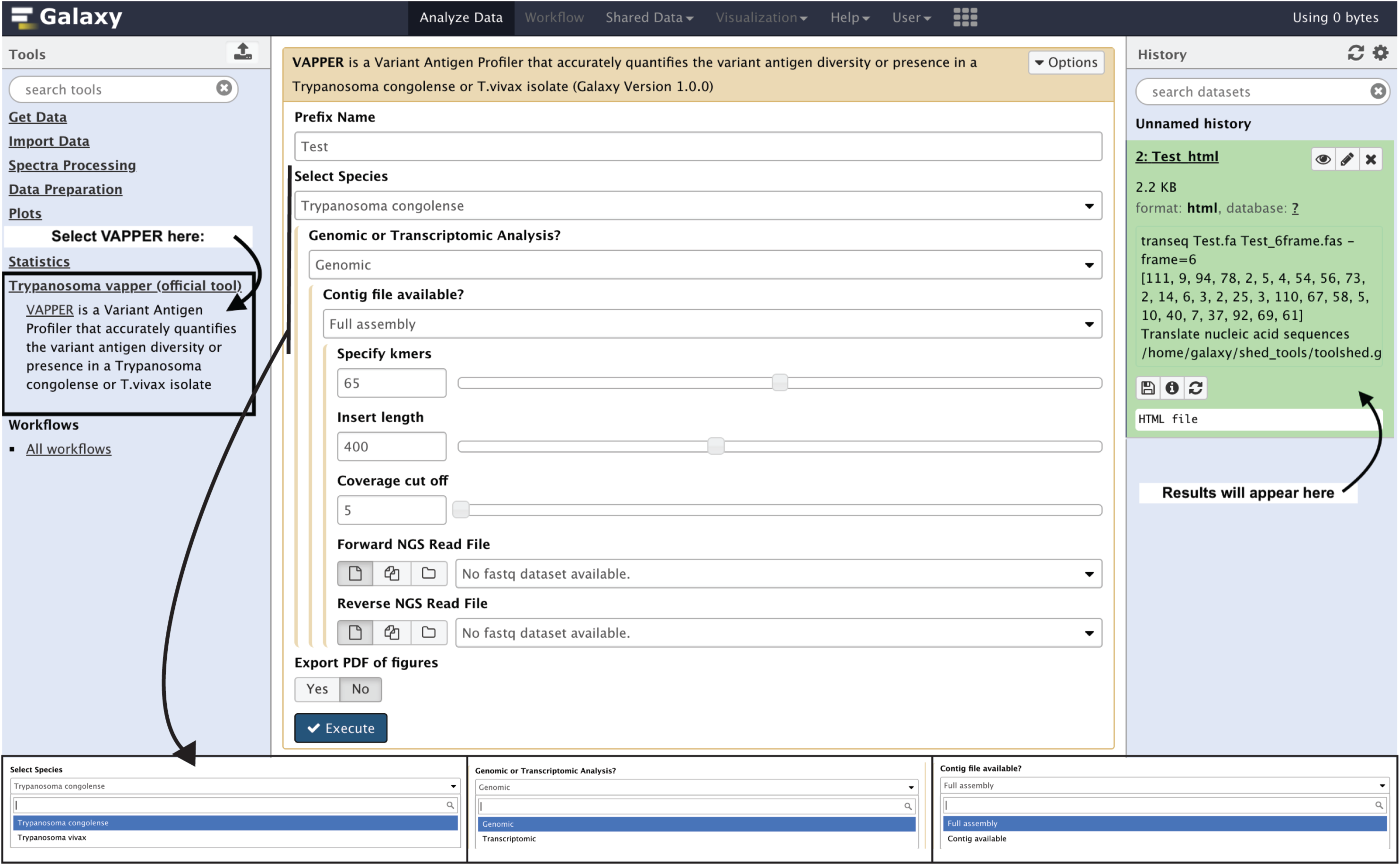
Screenshot of VAPPER on the Galaxy interface. This interface is available after installation of VAPPER from the Galaxy ToolShed [19] into a local Galaxy server. In this case, VAPPER was installed on the University of Liverpool Galaxy server. The blue panel on the right shows how to search and select VAPPER after installation. The white panel at the centre shows the options available for the user, including the prefix name of the sample to appear on the output figures, the species, and the type of input data. If any genomic pipeline is selected, further options for genome assembly parameters are available. Finally, the user can choose whether to get the graphs in PDF format (default is PNG only).

For *T. vivax*, the gVAP is based on presence or absence of pre-defined *VSG* genes, rather than phylotype frequencies as described for *T. congolense*. The *T. vivax VSG* repertoire is composed of distantly related lineages with no evidence for recombination [14]. Therefore, unlike *T. congolense* and *T. brucei, VSG* genes are often conserved across multiple strains, allowing us to build a *VSG* database for the entire species. No *T. vivax* tVAP is currently offered due to the lack of enough transcriptomic data available for benchmarking, but work is on-going to add this function. VSG-containing contigs are identified using BLAST 2.7.1 to detect sequence homology with a *T. vivax VSG* database. This information is added to a regularly updated presence/absence binary matrix of *T. vivax* genomes (N=29) and applied to a Euclidean distance-based clustering analysis. The results are presented as a heatmap and dendrogram, putting the sample in the context of the remaining *T. vivax* genomes and their known countries of origin (Fig. 3).

**Figure 3.**
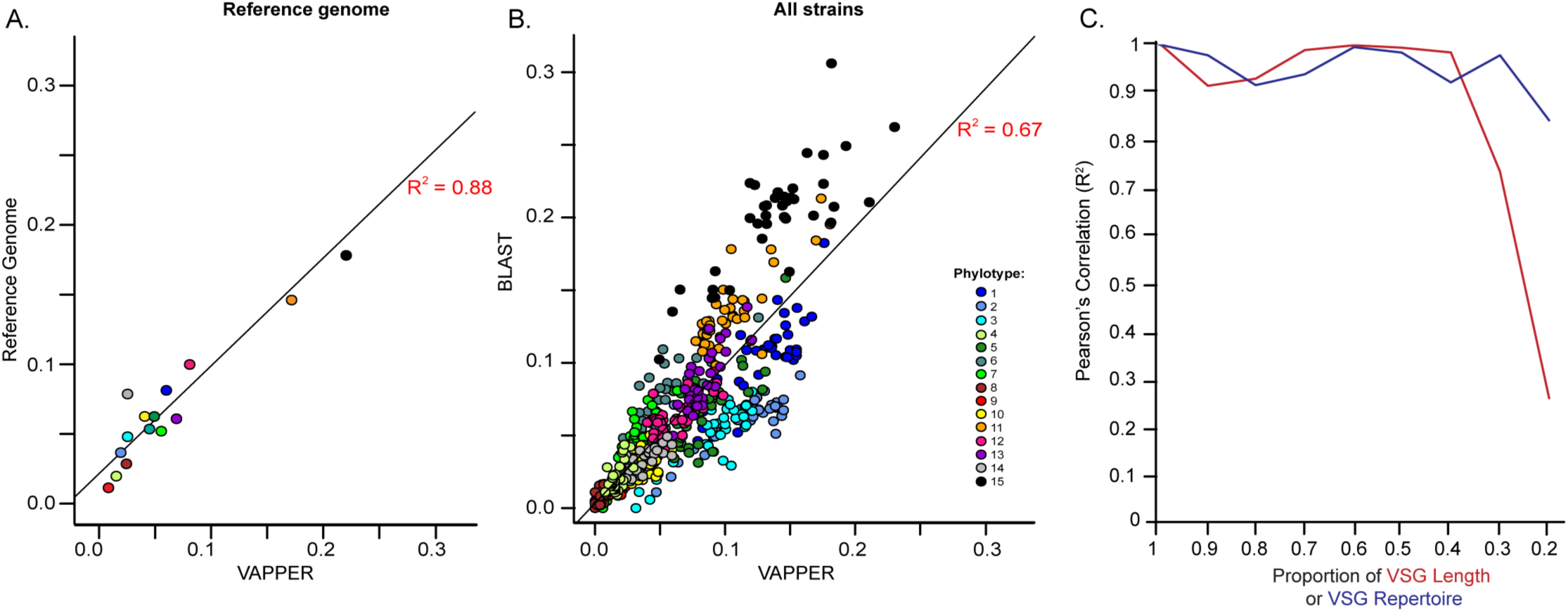
VAPPER performance (*T. congolense* genomic pipeline). (A) Correlation of phylotype frequencies produced by VAPPER and those manually curated in the *T. congolense* IL3000 reference genome sequence [14]. Pearson’s product moment correlation statistics: R^2^ = 0.88, t(13) = 9.7321, P < 0.001. (B) Correlation of phylotype frequencies produced by VAPPER and BLAST-based [41] phylotype frequencies in a panel of 41 *T. congolense* strains. Pearson’s product moment correlation: R^2^ = 0.64, t(566) = 34.39, P < 0.001. Phylotypes are color-coded according to the key. (C) VAPPER accuracy in fragmented (red) or incomplete (blue) genomes. Line graphs show correlations of the expected antigen profiles of a known set of VSGs sequences from the IL3000 genome sequence with antigen profiles produced from fragmented VSGs or incomplete VSG repertoires. Fragmentation and genome incompletion were simulated from random sampling. Gene fragmentation was calculated as a proportion of the mean length of the original VSG sequences (Mean±σ=1163±129 nucleotides). Figure adapted from [18].

In its Linux version, VAPPER can process multiple samples concurrently, providing that the input files are compiled in a single directory. Results are shown for all samples simultaneously, allowing direct comparison of variant antigen profiles across multiple isolates, conditions, or replicates. The tabular output can be incorporated in downstream statistical analysis, whilst the graphical outputs provide figures for the visualization of antigen repertoire variability.

### Linux Package Installation

To facilitate usage, the installation of VAPPER and its dependencies is automated. Upon first download of the software, a single script will ensure the system has all the required dependencies and install them in a local directory if necessary. In naïve environments and for users without administrator rights to install the necessary libraries, a Python virtual environment can be set upon each new session.

### The Galaxy Tool

VAPPER is available for installation in local Galaxy servers from the Galaxy ToolShed (https://toolshed.g2.bx.psu.edu/repository?repository_id=08b5616f1d3df20c). The purpose of the incorporation of VAPPER in Galaxy is to provide a simple front-end component for non-experienced users (Fig. 2). Results can be visualised directly in Galaxy, or can be downloaded as a compressed folder containing an HTML file with combined results, individual PNG and PDF files of the heatmaps, PCA plots, and bar charts produced, and the CSV files containing the raw values of phylotype proportions and deviation from the mean.

### Benchmarking

The performance of the *T. congolense* gVAP pipeline was compared to the manually annotated VAP of the IL3000 reference genome (Fig. 3A) and to the BLAST-based VAPs of 41 isolates (Fig. 3B) [18]. There is a very good correlation between profiles produced by VAPPER and the known IL3000 VAP (R^2^ = 0.88, t(13) = 9.7321, P < 0.001) and a good correlation with the BLAST-based method (R^2^=0.67, Pearson’s product moment correlation, t_(566)_=34.4, p < 0.001). Minor differences were further investigated and found to be due to BLAST’s difficulty in either analysing small contigs or quantifying multiple VSGs in the same contig sequence. Therefore, in general, more VSGs were recovered with VAPPER than with BLAST (Mean ± σ=721±277 vs. 669 ± 292, paired *t*-test, *p*-value = 0.005). A further strength of VAPPER is the ability to deal with poor, fragmented, genome assemblies. As described in our previous paper [18], when a single *VSG* gene is located in two distinct contig fragments, BLAST counts them incorrectly as separate genes, whereas VAPPER will not because the diagnostic motif is only present once. Therefore, we can now accurately calculate antigen profiles from incomplete genome assemblies (up to 30%), and with a *VSG* fragmentation level up to 40% of the original gene length (223 nucleotides) (Fig. 3C).

### Validation by example

#### *T. congolense* gVAP

We have used the VAPPER to analyse the genomic repertoire of 98 *T. congolense* samples of savannah and forest-subtypes, collected from 12 countries across Africa, and previously described by us [18] and others [25]. In Fig. 4, two heatmaps and corresponding dendrograms show how the VSG repertoires of each strain relate to each other. On the left, the heatmap represents phylotype proportion, i.e. how many genes a specific phylotype contains in the context of the complete *VSG* repertoire for a given strain (Fig. 4A). This heatmap shows that P4, 8, 9, 10, and 14 have few genes in all strains, whereas other phylotypes (e.g. P1, 2, 15) are more variable, being quite abundant in some strains and rare in others. The heatmap on the right shows phylotype deviation from the mean (Fig. 4B), which is calculated as the difference between the phylotype proportion shown in panel A and the arithmetic mean of phylotype proportions. The latter is calculated from the current database, thus it will change as new samples are added.

**Figure 4.**
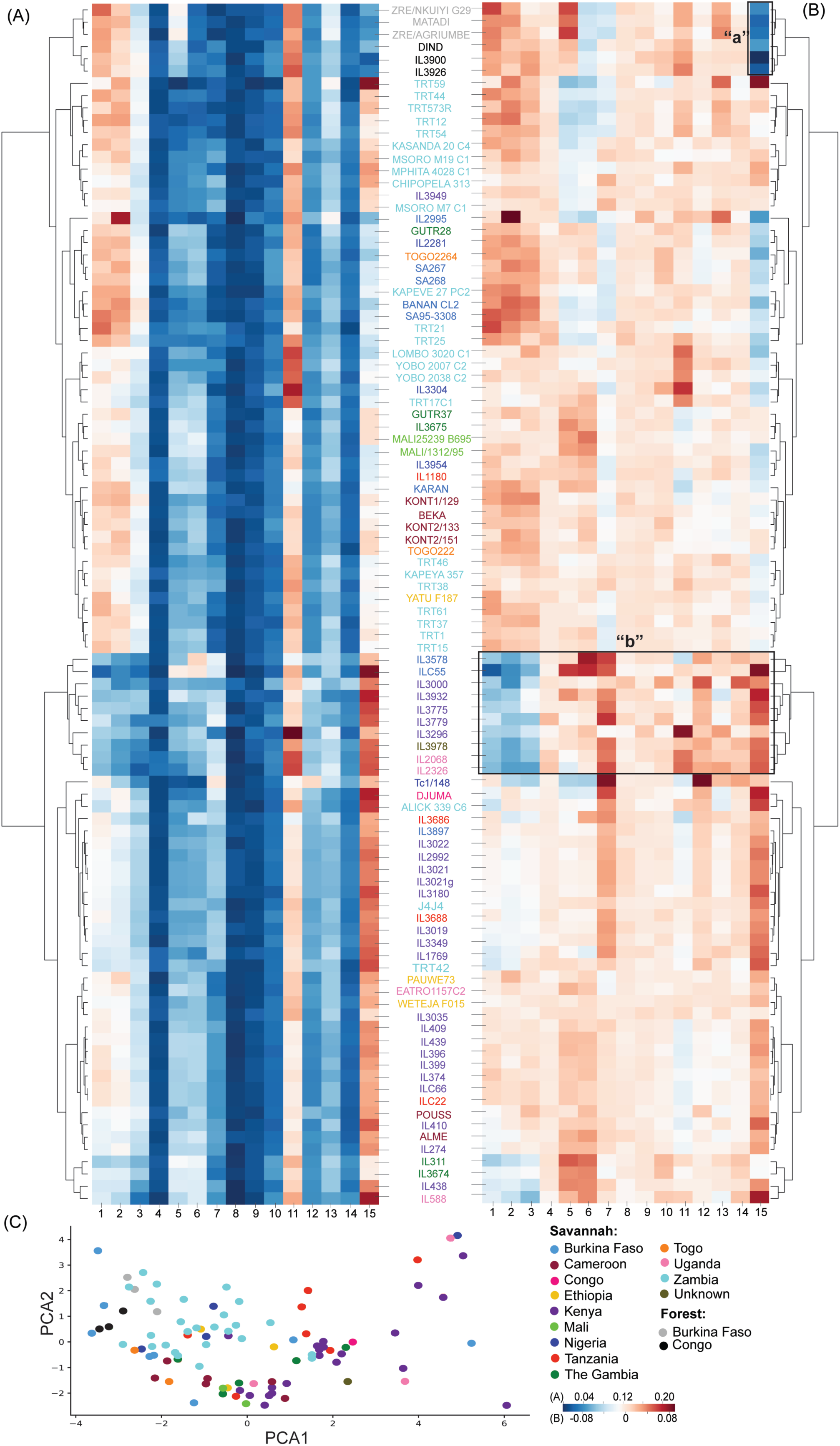
VAPPER output for *T. congolense* genomic pipeline. (A) Heatmap and corresponding dendrogram showing the variant antigen profiles (VAP) of the current genomic database expressed as phylotype frequencies [18, 25]. (B) Heatmap and corresponding dendrogram showing the variant antigen profiles (VAP) of the current genomic database expressed as deviation from the mean phylotype frequency [18, 25]. Labels “a” and “b” are referred to in the text. (C) PCA plot representing variation in *VSG* repertoire across the *T. congolense* genomic database [18, 25] (N=97).

The phylotype proportion variation patterns are perhaps better detected in the normalised heatmap (Fig. 4B). For example, it is possible to detect a signature of underrepresented P15 characteristic of all forest-subtype samples (denoted by “a”), abundant P15 in all Kenyan isolates (in purple), as well as a distinct pattern characteristic of strains IL3578 to IL2326, characterised by the combination of low P1 to 3 and high P7 (denoted by “b”). The latter does not seem to be related to geography, as it encompasses isolates from Kenya, Uganda, and Burkina Faso. The PCA plot further indicates that VSG repertoires and geography are only weakly correlated (Fig. 4C), which agrees with our previous observation that *T. congolense* VSG repertoires do not mimic either population structure or geography [18].

#### *T. congolense* tVAPs

We have used VAPPER to analyse the expressed VSG repertoire of the metacyclic (infective) life stage of *T. congolense*. For that, we have produced a tVAP for the strain TC13, whose transcriptome was published by Awuoche *et al*. (2018) [31]. We have compared its metacyclic tVAP to the metacyclic tVAP of the 1/148 strain (MBOI/NG/60/1-148) that we have previously described [29]. Furthermore, we have compared them to the genomic *VSG* repertoires of the same strain, or a related one (Fig. 5). As we do not have a genome sequence for the TC13 isolate, we compared it to IL3000, which was isolated in the same region (Transmara, Kenya) [32].

**Figure 5.**
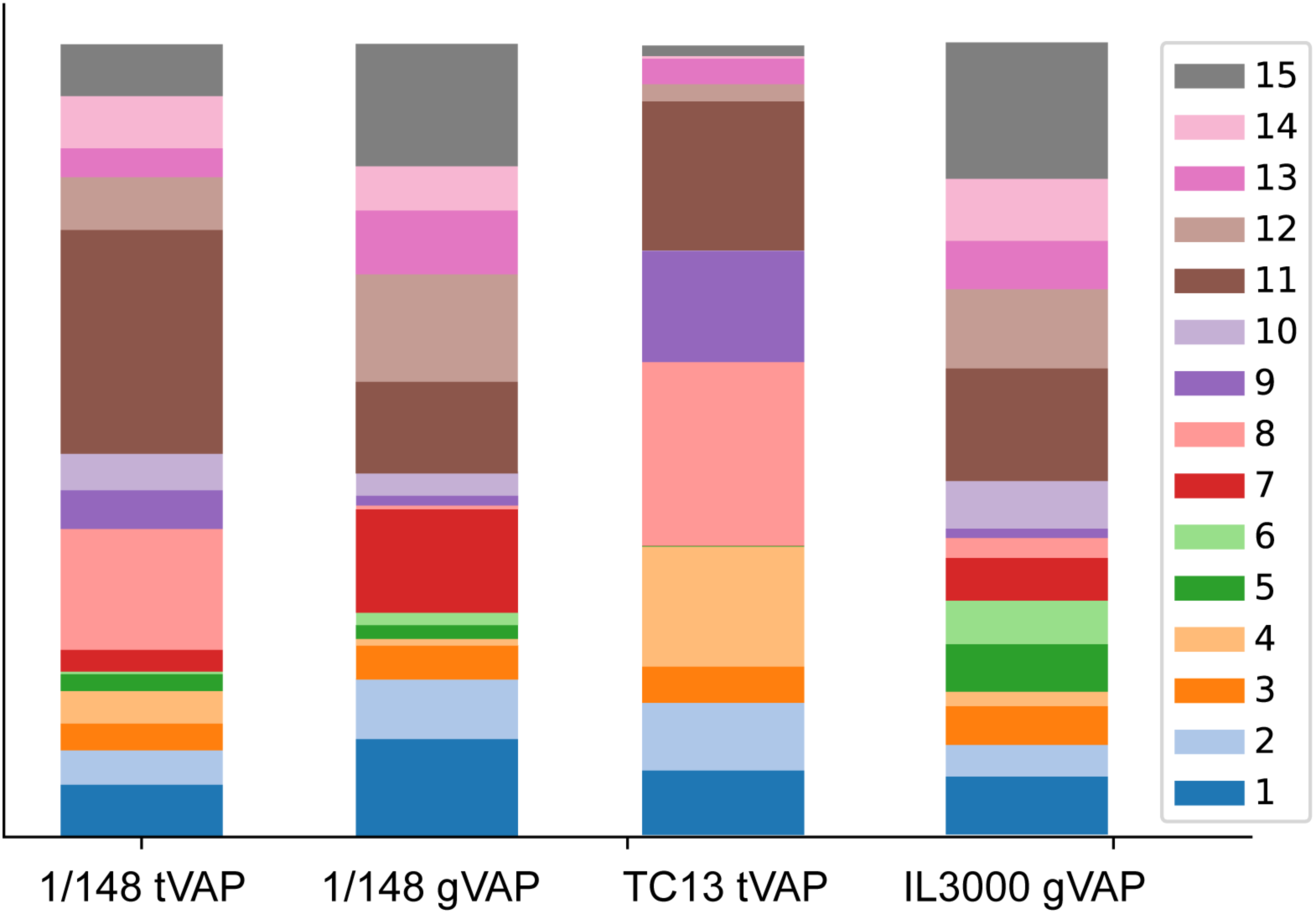
VAPPER output for *T. congolense* transcriptomic pipeline. Stacked bar charts showing expressed variant antigen profiles (VAPs) of metacyclic-stage *T. congolense* from strain 1/148 [18] and TC13 [31] compared to the genomic repertoires of the same strain (1/148) or a closely related one (IL3000) [28]. Phylotypes are colour-coded according to key. Size of each stack represents proportion of the phylotype relative to the total repertoire of expressed *VSGs*.

When we compare the gVAPs of 1/148 and IL3000, we see that they are distinct, and so are the tVAPs (e.g. P4 is more represented in TC13, whereas P10 is more represented in 1/148 than in TC13). However, P8 is overrepresented in both isolates compared to the genomic repertoires (Fig. 5). This agrees with our previous observation that the pattern of metacyclic *VSG* expression is significantly different from the genome repertoires, and that the metacyclic *VSG* repertoire is particularly enriched for P8 genes [18]. With the analysis of the TC13 transcriptome, we can now add that this enrichment does not seem to be strain-specific, but rather equally applicable to *T. congolense* strains of distinct backgrounds.

#### *T. vivax* gVAP

The *T. vivax* gVAP shows the VAPs in the context of the sample cohort (N=29), which currently includes samples from Nigeria, Uganda, Gambia, Ivory Coast, Brazil, Burkina Faso, and Togo. The dendrogram represents the relationships between the multiple strains, whereas the heatmap shows whether *VSG* genes are present or absent in each strain (Fig. 6A). The VAP relationship shows a separation between Nigerian (in dark blue) and the remaining samples, as well as a clear difference between Brazilian and Ugandan isolates. The geographical signature is diminished slightly in the non-Nigerian West African strains, although this is may reflect the smaller number of samples per country and perhaps the geographical closeness between Togo, Burkina Faso, and Ivory Coast. Despite the lack of a transcriptomic pipeline for *T. vivax*, we can use the gVAP to understand the geographical distribution of expressed VSGs. As an example, we took the two most abundant *VSGs* in the transcriptomes of three strains (i.e. LIEM-176 from Venezuela [33], IL1392 from Nigeria [34], and Lins from Brazil [35]) and compared them to the VAP database (Fig. 6B). We observe that there are five different VSGs, which represent three different geographical patterns (Fig. 6C). Specifically, the first LIEM-176 *VSG* transcript has been found in strains from Venezuela, Nigeria and Gambia, but not in Brazil, Uganda, or Ivory Coast (map 1 in Fig. 6C). The second LIEM-176 *VSG* is present in Brazil, Venezuela, Nigeria, and Uganda, yet not in Ivory Coast, a pattern that is shared with the top two most abundant *VSGs* in Lins (map 2 in Fig. 6C). Finally, the top two most abundant *VSGs* in IL1392 have been found in strains from Brazil, Venezuela, Gambia, Nigeria, but not in Uganda nor Ivory Coast (map 3 in Fig. 6C). It is possible that strain or location-specific *VSGs* might be epidemiologically relevant, perhaps contributing to the considerable phenotypic variation observed in *T. vivax* AAT.

**Figure 6.**
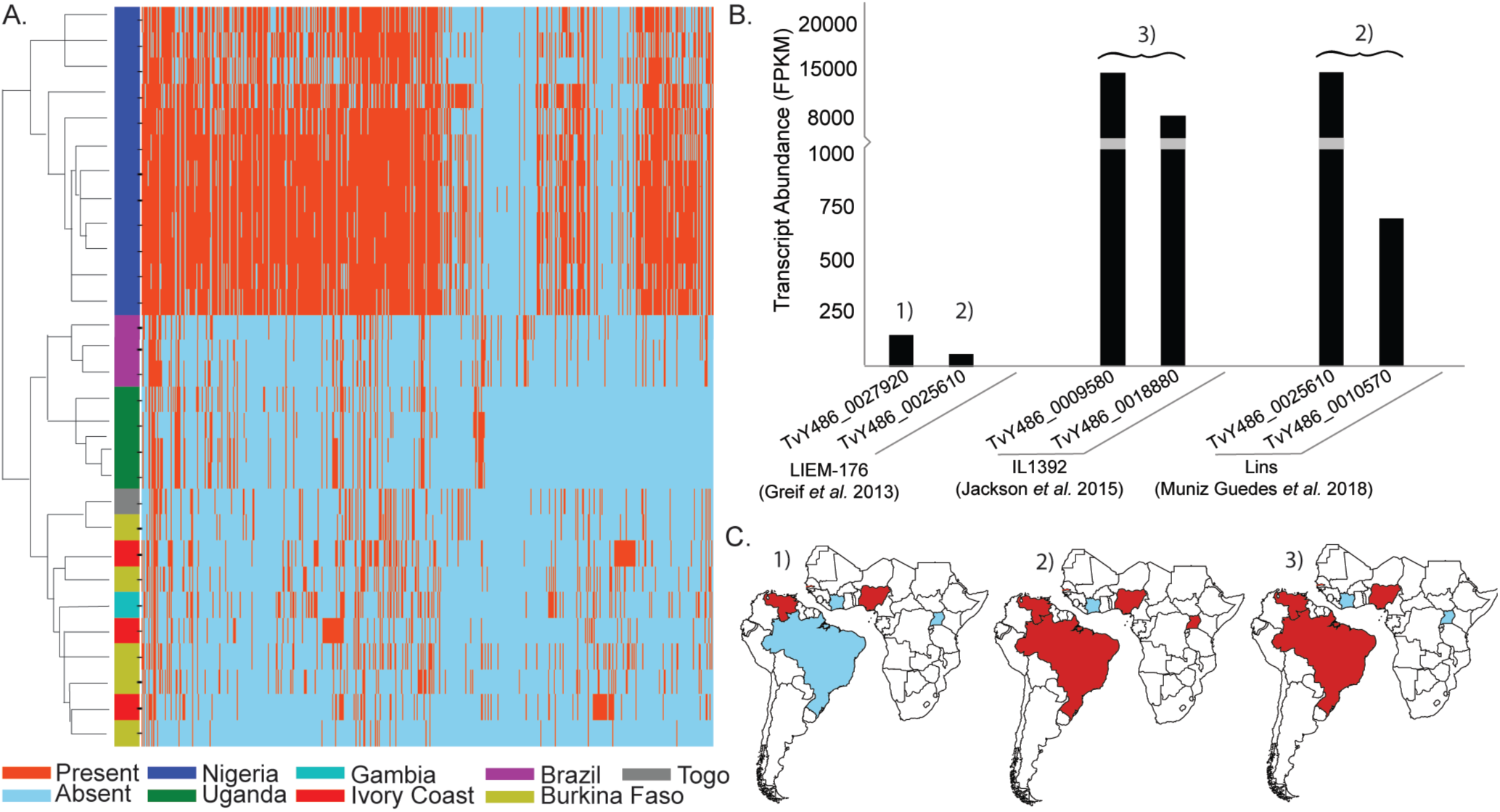
VAPPER output for *T. vivax* and its uses. (A) Heatmap and corresponding dendrogram showing the *T. vivax* variant antigen profiles (VAPs) in the context of the current genomic database (N=29). Strains are colour-coded according to key. (B) Two most abundant *VSG* genes in the three transcriptomes previously published for *T. vivax* (strains LIEM-176 [33], IL1392 [34], and Lins [35]. Numbers 1) to 3) relate to the *VSG* type in C. (C) Geographical distribution of the 6 *VSG* transcripts described in B. This is information can be obtained from the analysis of the VAP heatmap presented in (A).

## Conclusion

VAPPER is the first tool for the systematic analysis of *VSG* gene and expression diversity across strains and during infections. It establishes a practical approach for measuring antigenic diversity in these important pathogens based on universal protein motifs and/or gene mapping. Despite being often seen as a veterinary extension of HAT, AAT is a spectrum of diseases, dependent on the multiple species and strains of African trypanosomes and their multiple mammal hosts [36]. This predicament results in large variability in pathogenesis, epidemiology, and clinical outcome that remains poorly understood. For example, in East Africa, *T. vivax*, usually causes mild, chronic disease, but has occasionally resulted in acute haemorrhagic syndromes [37] without apparent reason. Likewise, in Brazil, related strains of *T. vivax* can cause both chronic disease of low parasitaemia and localized epidemics of up to 70% mortality rates, even in the same host species (although perhaps not the same genetic background) [38–40]. VAPPER allows us to identify and characterise differences in antigenic repertoires between strains, hosts, and conditions, which may be the starting point to build a real understanding of the association between disease genotypes and phenotypes. Importantly, with time this approach may be extended to the analysis of similar multi-copy, contingency gene families, particularly those involved in antigenic variation, in diverse pathogens.

## Availability and requirements

Project name: VAPPER – High-throughput Variant Antigen Profiling in African trypanosomes

Project home page: https://github.com/PGB-LIV/VAPPER

Operating System: Platform independent

Programming language: Python

Installation Requirements: Velvet 1.2.10; HMMER 3.1b2; Bowtie 2 2.2.6; SAMtools 1.6; Cufflinks 2.2.1; BLAST 2.7.1; EMBOSS

License: Apache v.2.0

## Declarations

Ethics approval and consent to participate

Not applicable.

## Consent for publication

Not applicable.

## Authors’ contributions

SSP wrote the original code in Perl and tested the software. JH and ARJ wrote the final code in Python. SSP and APJ conceptualized the software and wrote the manuscript. All authors contributed to and approved the final manuscript.

## References

1. Barry JD, Ginger ML, Burton P, McCulloch R. Why are parasite contingency genes often associated with telomeres? Int. J. Parasitol. 2003;33:29–45.

2. de la Fuente J, Lew A, Lutz H, Meli ML, Hofmann-Lehmann R, Shkap V, et al. Genetic diversity of anaplasma species major surface proteins and implications for anaplasmosis serodiagnosis and vaccine development. Anim. Health Res. Rev. 2005;6:75–89.

3. Kyes SA, Kraemer SM, Smith JD. Antigenic variation in Plasmodium falciparum: Gene organization and regulation of the var multigene family. Eukaryot. Cell. 2007;6:1511–20.

4. McCulloch R, Cobbold CA, Figueiredo L, Jackson A, Morrison LJ, Mugnier MR, et al. Emerging challenges in understanding trypanosome antigenic variation. Emerg. Top. Life Sci. 2017;1:585–92.

5. Pays E. The variant surface glycoprotein as a tool for adaptation in African trypanosomes. Microbes Infect. 2006;8:930–7.

6. Cross G a M, Kim HS, Wickstead B. Capturing the variant surface glycoprotein repertoire (the VSGnome) of Trypanosoma brucei Lister 427. Mol. Biochem. Parasitol. Elsevier B.V.; 2014;195:59–73.

7. Matthews KR, McCulloch R, Morrison LJ. The within-host dynamics of African trypanosome infections. Philos. Trans. R. Soc. Lond. B. Biol. Sci. 2015;370:20140288–.

8. Capewell P, Clucas C, DeJesus E, Kieft R, Hajduk S, Veitch N, et al. The TgsGP gene is essential for resistance to human serum in Trypanosoma brucei gambiense. PLoS Pathog. 2013;9:e1003686.

9. Uzureau P, Uzureau S, Lecordier L, Fontaine F, Tebabi P, Homblé F, et al. Mechanism of Trypanosoma brucei gambiense resistance to human serum. Nature. 2013;501:430–4.

10. De Greef C, Hamers R. The serum resistance-associated (SRA) gene of Trypanosoma brucei rhodesiense encodes a variant surface glycoprotein-like protein. Mol. Biochem. Parasitol. 1994;68:277–84.

11. Van Xong H, Vanhamme L, Chamekh M, Chimfwembe CE, Van Den Abbeele J, Pays A, et al. A VSG expression site-associated gene confers resistance to human serum in Trypanosoma rhodesiense. Cell. 1998;95:839–46.

12. Wiedemar N, Graf FE, Zwyer M, Ndomba E, Kunz Renggli C, Cal M, et al. Beyond immune escape: a variant surface glycoprotein causes suramin resistance in Trypanosoma brucei. Mol. Microbiol. 2018;107:57–67.

13. Salmon D, Geuskens M, Hanocq F, Hanocq-Quertier J, Nolan D, Ruben L, et al. A novel heterodimeric transferrin receptor encoded by a pair of VSG expression site-associated genes in T. brucei. Cell. 1994;78:75–86.

14. Jackson AP, Berry A, Aslett M, Allison HC, Burton P, Vavrova-Anderson J, et al. Antigenic diversity is generated by distinct evolutionary mechanisms in African trypanosome species. Proc. Natl. Acad. Sci. U. S. A. 2012;109:3416–21.

15. Marcello L, Menon S, Ward P, Wilkes JM, Jones NG, Carrington M, et al. VSGdb: A database for trypanosome variant surface glycoproteins, a large and diverse family of coiled coil proteins. BMC Bioinformatics. 2007;8:1–8.

16. Weirather JL, Wilson ME, Donelson JE. Mapping of VSG similarities in Trypanosoma brucei. Mol. Biochem. Parasitol. 2012;181:141–52.

17. Mugnier MR, Cross GAM, Papavasiliou FN. The in vivo dynamics of antigenic variation in Trypanosoma brucei. Science (80-.). 2015;347:1470–3.

18. Silva Pereira S, Casas-Sanchez A, Haines LR, Absolomon K, Ogugo M, Sanders M, et al. Variant antigen repertoires in Trypanosoma congolense populations and experimental infections can be profiled from deep sequence data with a set of universal protein motifs. Genome Res. 2018;28:1383–94.

19. Blankenberg D, Von Kuster G, Bouvier E, Baker D, Afgan E, Stoler N, et al. Dissemination of scientific software with Galaxy ToolShed. Genome Biol. 2014.

20. Afgan E, Baker D, van den Beek M, Blankenberg D, Bouvier D, Čech M, et al. The Galaxy platform for accessible, reproducible and collaborative biomedical analyses: 2016 update. Nucleic Acids Res. 2016;44:W3–10.

21. Cock PJA, Fields CJ, Goto N, Heuer ML, Rice PM. The Sanger FASTQ file format for sequences with quality scores, and the Solexa/Illumina FASTQ variants. Nucleic Acids Res. 2009;38:1767–71.

22. Pearson WR, Lipman DJ. Improved tools for biological sequence comparison. Proc. Natl. Acad. Sci. 1988;85:2444–8.

23. Zerbino DR. Using the Velvet de novo assembler for short-read sequencing technologies. Curr. Protoc. Bioinforma. 2010.

24. Eddy SR. A new generation of homology search tools based on probabilistic inference. Genome Inform. 2009;23:205–11.

25. Tihon E, Imamura H, Dujardin J-C, Van Den Abbeele J, Van den Broeck F. Discovery and genomic analyses of hybridization between divergent lineages of Trypanosoma congolense, causative agent of Animal African Trypanosomiasis. Mol. Ecol. 2017;

26. Langmead B, Salzberg SL. Fast gapped-read alignment with Bowtie 2. Nat Methods. 2012;9:357–9.

27. Trapnell C, Roberts A, Goff L, Pertea G, Kim D, Kelley DR, et al. Differential gene and transcript expression analysis of RNA-seq experiments with TopHat and Cufflinks. Nat. Protoc. 2012;7:562–78.

28. Gibson W. The origins of the trypanosome genome strains Trypanosoma brucei brucei TREU 927, T. b. gambiense DAL 972, T. vivax Y486 and T. congolense IL3000. Parasit. Vectors. BioMed Central Ltd; 2012;5:71.

29. Young CJ, Godfrey DG. Enzyme polymorphism and the distribution of Trypanosoma congolense isolates. Ann. Trop. Med. Parasitol. 1983;77:467–81.

30. Abbas AH, Pereira SS, D’Archivio S, Wickstead B, Morrison LJ, Hall N, et al. The structure of a conserved telomeric region associated with variant antigen loci in the blood parasite Trypanosoma congolense. Genome Biol. Evol. 2018;evy186.

31. Awuoche EO, Weiss BL, Mireji PO, Vigneron A, Nyambega B, Murilla G, et al. Expression profiling of Trypanosoma congolense genes during development in the tsetse fly vector Glossina morsitans morsitans. Parasit. Vectors. Parasites & Vectors; 2018;11:1–18.

32. Ferrante A, Allison AC. Alternative pathway activation of complement by African trypanosomes lacking a glycoprotein coat. Parasite Immunol. 1983;5:491–8.

33. Greif G, Ponce de Leon M, Lamolle G, Rodriguez M, Piñeyro D, Tavares-Marques LM, et al. Transcriptome analysis of the bloodstream stage from the parasite Trypanosoma vivax. BMC Genomics. 2013;14:149.

34. Jackson AP, Goyard S, Xia D, Foth BJ, Sanders M, Wastling JM, et al. Global Gene Expression Profiling through the Complete Life Cycle of Trypanosoma vivax. PLoS Negl. Trop. Dis. 2015;9:e0003975.

35. Guedes RLM, Rodrigues CMF, Coatnoan N, Cosson A, Cadioli FA, Garcia HA, et al. A comparative in silico linear B-cell epitope prediction and characterization for South American and African Trypanosoma vivax strains. Genomics. Elsevier; 2018;0–1.

36. Morrison LJ, Vezza L, Rowan T, Hope JC. Animal African Trypanosomiasis: Time to Increase Focus on Clinically Relevant Parasite and Host Species. Trends Parasitol. 2016;32:599–607.

37. Wellde BT, Chumo DA, Adoyo M, Kovatch RM, Mwongela GN, Opiyo EA. Haemorrhagic syndrome in cattle associated with trypanosoma vivax infection. Trop. Anim. Health Prod. 1983;15:95–102.

38. Paiva F, De Lemos R a a, Nakazato L, Mori a E, Brum KB, Bernardo KC, et al. Trypanosoma Vivax Em Bovinos No Pantanal Do Estado Do Mato Grosso Do Sul, Brasil□: I - Acompanhamento Clínico,. Rev. Bras. Parasitol. Veterinária. 2000;9:135–41.

39. Cadioli FA, de Athayde Barnabé P, Zacarias Machado R, Alves Teixeira MC, André MR, Sampaio PH, et al. First report of Trypanosoma vivax outbreak in dairy cattle in São Paulo state, Brazil. Rev. Bras. Parasitol. Vet., Jaboticabal. 2012;21:118–24.

40. Fidelis Jr OL, Sampaio PH, Machado RZ, Andre MR, Marques LC, Cadioli FA. Evaluation of clinical signs, parasitemia, hematologic and biochemical changes in cattle experimentally infected with Trypanosoma vivax. Brazilian J. Vet. Parasitol. 2016;2961:69–81.

41. Altschul SF, Gish W, Miller W, Myers EW, Lipman DJ. Basic local alignment search tool. J. Mol. Biol. 1990;215:403–10.

